# A parametric interpretation of Bayesian Nonparametric Inference from Gene Genealogies: linking ecological, population genetics and evolutionary processes

**DOI:** 10.1101/200469

**Authors:** José M. Ponciano

## Abstract

Using a nonparametric Bayesian approach Palacios and Minin [1] dramatically improved the accuracy, precision of Bayesian inference of population size trajectories from gene genealogies. These authors proposed an extension of a Gaussian Process (GP) nonparametric inferential method for the intensity function of non-homogeneous Poisson processes. They found that not only the statistical properties of the estimators were improved with their method, but also, that key aspects of the demographic histories were recovered. The authors’ work represents the first Bayesian nonparametric solution to this inferential problem because they specify a convenient prior belief without a particular functional form on the population trajectory. Their approach works so well and provides such a profound understanding of the biological process, that the question arises as to how truly “biology-free” their approach really is. Using well-known concepts of stochastic population dynamics, here I demonstrate that in fact, Palacios and Minin’s GP model can be cast as a parametric population growth model with density dependence and environmental stochasticity. Making this link between population genetics and stochastic population dynamics modeling provides novel insights into eliciting biologically meaningful priors for the trajectory of the effective population size. The results presented here also bring novel understanding of GP as models for the evolution of a trait. Thus, the ecological principles foundation of Palacios and Minin [1]’s prior adds to the conceptual and scientific value of these authors’inferential approach. I conclude this note by listing a series of insights brought about by this connection with Ecology.

## 1. Introduction

Statistical inference for stochastic processes in biology was central to the research in Paul Joyce’s lab. I was humbled and challenged by the request to write a paper celebrating the memory of Paul Joyce’s contributions to mathematical modeling and statistical inference in population genetics. Little by little, my fears became excitement when I envisioned a little note illustrating the type of interactions that would occur between the members of his lab, and anyone who approached him to talk about science. Those interactions often occurred very early in the morning, around seven AM, his favorite moment of the day to indulge in research (with coffee).

At the time I started to study under his guidance (summer 2002), professors Zaid Abdo and Vladimir Minin were my lab mates. I had the privilege to learn with and from them through day to day conversations, classes, homeworks, research problems and most importantly, from our successes and failures. The diversity of topics that we talked about and worked on was naturally, a reflection of Paul’s innate fascination for any problem in biology having to do with mathematical statistics and stochastic processes. Indeed, he would often be the glue connecting the thinking and ideas among topics. Seeking to see beyond a particular area or application, and understand the connections between probabilistic results applied to one or another area in biology is perhaps, one of the most valuable lessons I got from him.

During one of my last visits to Idaho before his tragic accident, we reminisced about the times when “Vlad” (Minin) was a student. We naturally talked about his (then) latest work, a successful attempt to dramatically improve the accuracy and precision of Bayesian inference of population size trajectories from gene genealogies [1]. During the rest of our conversation, I proceeded to build a case to demonstrate why I thought that this novel methodology had a remarkable ability to recapitulate fundamental biological properties of the system: because unbeknownst to them, Palacios and Minin’s contribution was strongly connected with theoretical concepts and results from statistical ecology. My argument met, of course, a skeptic listener but after my exposition and many interjections, Paul apparently conceded because he exclaimed: “Well I hope you’re right, because if so, then this would be one of these cool instances in which mathematical population genetics learns from ecological thinking”. The reasoning I presented to Paul, formally written, constitutes the contents of this note.

Palacios and Minin [1] proposed a Bayesian nonparametric methodology to reconstruct past population dynamics using genomic data and the Coalescent process. This non homogeneous Markov death process specifies the relationship between ancestral genealogies of a random sample of genes and effective population size. Because changes in population size result in changes on the genetic pool in a population, at any point in time genomic data carries information regarding past demographic processes and population dynamics. Although estimating the effective population size amounts to estimating the total population size in an idealized Wright-Fisher model, studying changes in this parameter remains important because of its interpretation as a metric of relative genetic diversity.

Motivated by the lack of statistical methods to infer past population dynamics from a sample of genes that didn’t depend on strong parametric assumptions, Palacios and Minin [1] proposed a transformed Gaussian Process (GP) as the prior for past population trajectories. These authors justify their choice because such process “does not adhere to a particular functional form, or hypothesis on past population dynamics” [1]. In this article, I borrow results from theoretical ecology, to show that Palacios and Minin [1] prior choice, although justifiable under numerical and statistical grounds, can be interpreted as a class of stochastic population dynamics models, albeit one previously not studied and hence, one that brings novel insights into both population genetics and statistical ecology.

Engen et al. [2] published what now is considered one of the standard references to understand the concepts of “demographic stochasticity” and “environmental variability (stochasticity)” in population dynamics modeling. These authors drew their ideas from the stochastic processes models of Keiding [3] and Ludwig [4] which incorporated two main sources of stochasticity: stochasticity due to random births and deaths, known as demographic stochasticity; and temporal stochasticity in any of the demographic rates (*e.g.* good years/bad years for survival, etc…). Traditional ecological concepts, such as density-dependence (the regulation of population growth rates according to the density of such population) were also explicitly incorporated in these models. Operationally, formulating a model with the so called ‘demographic stochasticity’ amounted to specify, for instance, a Branching Process (BP) model with a density dependent offspring distribution of individuals. To add temporal stochasticity into one of the demographic rates, or what came to be known as ‘environmental stochasticity’ [5], a temporally uncorrelated random shock was added to the mean of the offspring distribution (often assumed to be Poisson). The result was a density-dependent, BP in Random Environments (BPRE) model [6]. At that time, various properties of simpler BPRE’s had already been worked out by Athreya and Karlin [7, 8].

Diffusion approximations of the BPRE models later opened the door to the study of animal abundance fluctuations as modeled by realistic, stochastic population dynamics models [3, 4, 6, see Appendix 1]. Straightforward analytical expressions of the properties of the density-dependent BPRE models (such as stopping times and quasiextinction probabilities) are often too unwieldy or difficult to obtain. Their approximation by means of diffusion processes however, have led to a remarkable improvement in the understanding of how stochasticity from demographic events (births, deaths, etc.) and hence persistence, are affected when the rates themselves are allowed to vary randomly over time. To date, research in this field has yielded a plethora of results that guide the decisions and questions of wildlife managers, population biologists and theoretical ecologists alike [9, 10, 11, 12, 13, 14, 15, 16, 17, 18, 19, 20, 21, 22, 23, 24].

The diffusion approximation of ecological BP models are usually presented as a Stochastic Differential Equation (SDE) model [6]. The infinitesimal mean of these models usually corresponds to one of the well-known deterministic ODE models of population growth, such as the logistic equation. If only demographic stochasticity is considered (*i.e.*, if a BP model in constant environments is approximated with a diffusion), then the infinitesimal variance of the process scales proportionally to population size, whereas including both environmental and demographic stochasticities results in an infinitesimal variance with two terms, one proportional to population size and one that scales like the square of population size [see 25, and citations therein]. Finally, a density-dependent (or density-independent) SDE model of population abundances where the infinitesimal variance scales only like the square of population size has been shown to correspond to a model that assumes no demographic stochasticity and only environmental stochasticity. In what follows, first I briefly summarize the approximation of BPRE’s with diffusions. I then expose the relationship between Palacios and 95 Minin [1]’s prior for the effective population size and stochastic demography. I conclude by showing how, unbeknownst to Palacios and Minin [1], their GP model brings about a novel parametric understanding of stochastic population dynamics.

## 2. Palacios and Minin’s model and Stochastic Demography

At the core of these author’s approximation is the usage of a transformation of a GP as a prior for the effective population size, *N*_*e*_(*t*). GP are stochastic processes such that any finite sample from the process has a joint multivariate normal distribution [26]. As I explain below, this defining property of GPs is crucial for Bayesian inference of a quantity that varies through time, like *N*_*e*_(*t*).

In the context of Bayesian statistics, the ‘nonparametrics’ labeling refers to placing priors to a potentially infinite number of parameters. This approach differs from the classic definition of nonparametric (*e.g.* distribution free) statistics. Palacios and Minin’s inference is nonparametric in the sense that they do not adopt any particular functional form for past changes in effective population size (like exponential or logistic growth back from past to present). Their contribution is novel, because instead of choosing from a set of prior beliefs consisting of different functional forms of time for these changes, the authors chose to model the prior for the past trajectory of the effective population size as a collection of points all drawn at random from a general stochastic process. This stochastic process then becomes the prior for the parameter of interest: the entire trajectory of the effective population size *N*_*e*_(*t*). As Rasmussen and Williams [26] put it, a function of time *f*(*t*) can be loosely thought of as a very long vector where each entry in the vector specifies the function value *f*(*t*) at a particular time *t* [26]. In Bayesian Inference, the difficulty imposed by having to specify an infinite dimensional object like a function of time as a prior is nicely overcome with GPs. Because finite samples from GPs are jointly multivariate normal, eliciting a prior for the function of interest at a finite number of points in time (here at a collection of points of *N*_*e*_(*t*)) loosely amounts to sampling from a multivariate normal distribution at those points. As Rasmussen and Williams [26] explain, the resulting inference gives the same answer as if the infinitely many other points were taken into account. In this particular case, following Adams et al [27], Palacios and Minin go one step beyond and use a transformation of a GP to elicit a flexible prior for the changing effective population size back in time, *N*_*e*_(*t*).

Flexibility of a GP prior is obtained by tuning the general properties of the process. These properties are completely specified by the GP mean and covariance functions. Changing these two functions results in the specification of different GP models. One of the best known GP models is the Ornstein Uhlenbeck (OU) process. As a GP, its values at multiple points in time have a joint multivariate normal distribution. Furthermore, sampled at regular discrete time intervals the OU process is an Autoregressive process of order 1 (an AR(1) process), a model well-kown in the field of time series analysis [22]. In particular, if at the initial time point 0, the OU process *X*_0_ is *x*_0_, then the conditional distribution of the process at any other point *t* is normal with mean and variance given by [22, 28]

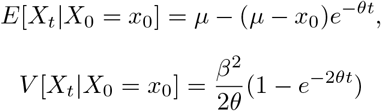

where *µ*, *θ* and *β*^2^ are the parameters that control the process. The covariance elements of the multivariate normal process at two different times *t*, *s* > 0 is given by

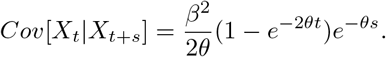

As time grows large, the distribution of the process attains a normal stationary distribution with mean

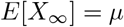

and variance

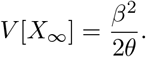

Now, in their usage of a transformation of a GP as a prior for *N*_*e*_(*t*), Palacios and Minin were exploiting the fact that a general GP can be thought to represent a vague prior. If the information is the data is strong enough, the vagueness in such prior would be overcome by the steepness of the likelihood function. A subjective Bayesian however, would seek to formulate a prior that embodies a biological mechanism by adopting a particular GP covariance structure. Under a likelihood approach, specifying a biology-based GP model would be a way to propose a hierarchical, state-space model that would embody a particular scientific hypothesis.

The point of this paper is to hypothesize and demonstrate through a simple mathematical development, that Palacios and Minin’s approach works very well, perhaps because un-intentionally, the GP transformation they proposed as a prior for *N*_*e*_(*t*) is in fact non-other than a special stochastic population dynamics model albeit with very deep connections with the theory of population biology. By exploring those deep connections I show below that their prior is in fact the point of entry to a plethora of biologically motivated GP priors. Such view could motivate eliciting a vast array of novel hypotheses that would seek to understand the processes behind the temporal fluctuation in *N*_*e*_. Finally, it is important to note that those connections with population biology are possible because of the Stochastic Differential Equation (SDE) representation of GP models like the OU process. Written as such, the OU model is a type of stochastic process known as a diffusion process [29]. In what follows I briefly explore those connections. The mathematical details are presented in the Appendix.

### 2.1. Diffusion approximation of density-dependent BP in random environments

In this section I briefly summarize the work of numerous authors, which work under different hypotheses and notations [30, 3, 4, 31, 6, 32, 33, 34]. This account is accompanied by a detailed appendix and is the basis for a later discussion of the key assumptions of the diffusion approximation of a BPRE, and its relation to Palacios and Minin [1]’s GP-based model for *N*_*e*_(*t*). These author’s prior for *N*_*e*_(*t*) is given by the transformation

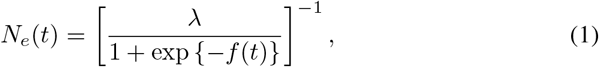

where *f*(*t*) is a Gaussian Process (GP). By so doing, the authors posit a priori sigmoidal, scaled logistic function of a GP with a restricted range between (0, λ) for 1/*N*_*e*_(*t*). Their approach to build a prior is then purportedly devoid of an explicit biological mechanism (*phenomenological*). For the sake of computational tractability, the authors use a Markovian GP and compare, for one of their examples, the performance of Brownian motion, an OU process and a Brownian Motion (BM) model. In the case of the BM and OU diffusion processes, because a smooth transformation of a diffusion process is also a diffusion [see 29, Theorem 2.1 p. 173], Palacios and Minin [1]’s prior for *N*_*e*_(*t*) is also a diffusion. The key point of this note is to show that the resulting diffusion prior for *N*_*e*_(*t*) is in fact a class of well known diffusion processes representing density-dependent, stochastic population growth. Hence, Palacios and Minin [1]’s prior for *N*_*e*_(*t*) is in fact, parametric, in the sense that can be made to represent different hypotheses regarding population growth, density dependence and the structure of stochasticity in a population.

Diffusions that model stochastic population growth are usually presented as SDEs of the form

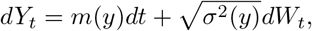

where *m*(*y*) is the infinitesimal mean of the diffusion, *σ*^2^(*y*) is its infinitesimal variance and *dW*_*t*_~ N(0, dt). Both *m*(*y*) and *σ*^2^(*y*) are continuous functions of *y*. In these models, the infinitesimal mean is usually given by the deterministic skeleton of an Ordinary Differential Equation (ODE), logistic-type model. The authors above have shown that a suitable diffusion approximation of a BP with a density dependent offspring distribution has an infinitesimal variance of the the form *σ*^2^(*y*) = *yα*, *α* > 0, whereas a diffusion approximating a BPRE has an infinitesimal variance of the form *σ*^2^(*y*) = *yα* + (*yβ*)^2^, β > 0. Thus, if the mean of the offspring distribution of a BP varies randomly every generation, then the infinitesimal variance of the corresponding approximating diffusion scales like the square of population size. This scaling then represents random fluctuations in the quality of the environment (*i.e.*, for instance, if iso there are good/bad years for reproduction), and is analogous to the variance scaling brought about by random, temporal changes in selection of population genetics models [29]. As I show in the following sections, the model for *N*_*e*_(*t*) that Palacios and Minin [1] present as a transformed GP is in fact, to a very close approximation, a stochastic logistic-type model with only environmental fluctuations.

In ecology, a plethora of sigmoidal mathematical models describing population growth as a function of continuous time have been reported in the literature [*e.g.* 35]. In light of empirical data, and among many equations in a large family of models related to Malthus’ “law of geometric growth”, it has been shown that the Gompertz equation emerges as one of the best models of the growth of population size as a function of time [see citations in 36, 22, 24]. In bacterial growth research for instance, the Gompertz model has served as a golden reference to which models that account for various idiosyncratic phenomena of microbial cultures have been compared [37]. The solution of the Gompertz ODE model *dy*_*t*_/*dt* = *θy*_*t*_[ln*κ* − ln*y*_*t*_], where κ is the carrying capacity and *θ* is the speed of equilibration is given by *y*_*t*_ = *κ* exp (*e*^−*θt*^ln(*y*_0_/*κ*)), where *y*_0_ is the initial population size. This solution has an inflection point at *κ*/*e*, provided *y*_0_ is below the carrying capacity. The stochastic Gompertz diffusion written in SDE form is given by [22]

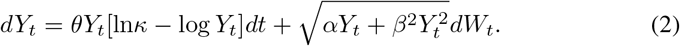

The construction of this diffusion starts with a family of BP’s denoted as 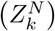, where *κ* = 0,1,2,…indexes the time and *N* represents some arbitrarily large population size that represents a key quantity in the model. It is a family of BP models because a different model is obtained with different values of *N*. This is akin to the formulation of the Wright-Fisher model with different (gene) population sizes. In the case of the stochastic Gompertz model derivation from these BP models, the *N* represents an unusually large population size. For example, [30, 3, 4, 31, 6] all use the carrying capacity of the logistic model as *N* (see explicit derivation in the appendix). The idea is then to accelerate time by *N* (*i.e.* so that *N* generations of the original process occur in one new unit of time) and scale the state by a factor 1/*N*. Writing the scaled process as 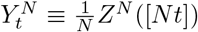, where [*Nt*] is the smallest integer close to *Nt*, then our diffusion *Y*_*t*_ is defined as 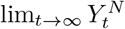. Now, such approximation works provided

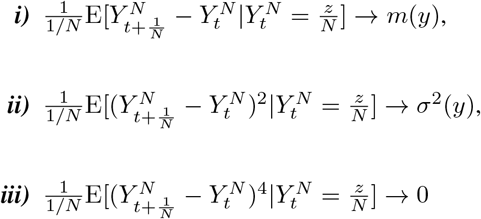

as *N* → ∞ and *z*/*N* → *y* [29], where *m*(*y*) and *σ*^2^(*y*) are continuous functions of *y*. To obtain the diffusion approximation, the moments of the one-step differences in the unscaled process 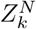 are first computed. Then, the process’ time unit and state are re-scaled, and this allows the calculation of the infinitesimal moments and checking if the limits above (conditions *i-iii*) hold. In the Appendix, I present as an example the steps leading to the construction of the stochastic Gompertz diffusion model with environmental and demographic stochasticity (eq. (2)). Although going through these calculations is a simple exercise, understanding this process is crucial to understand the rest of the ideas presented here, and in particular, to understand the population dynamics interpretation of Palacios and Minin [1]. But perhaps, the main reason why I kept this explanation here is that, while writing, I remembered Paul making fun of any of us (and of himself) whenever the word “Clearly…” appeared as the connection between two equations in a a manuscript or paper. In talks, he would seldom miss the opportunity to explain even the most elementary math steps. And one had to be vigilant, because those same explanations were often the key to understand the punchline of his message.

### 2.2. The Gompertz SDE and the OU process

Let *Y*_*t*_ be a Gompertz diffusion with environmental stochasticity and no demographic stochasticity [22]. Then, its infinitesimal mean and variance are given by *m*_*Y*_(*y*) = *θy*[ln κ – ln*y*] and 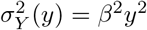 respectively. In SDE form we write

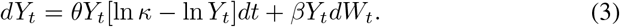

Let *X*_*t*_ = *g*(*Y*_*t*_) be a smooth invertible transformation of *Y*_*t*_. Then, it immediately follows that *X*_*t*_ is also a diffusion with infinitesimal mean and variance given by 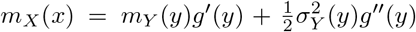 and 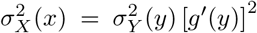 respectively [29], where *y* = *g*^−1^(*x*). This transformation of an SDE is known as Ito’s transformation or “It̂o’s formula” (see [28]), and differs from what one would obtain by using R. Stratanovich’s definition of stochastic integrals.[28, 32, 33] clarify such differences. All the stochastic integrals in this paper follow It̂o’s stochastic calculus. Then, it follows that setting *g*(*y*) = ln *y* yields *m*_*x*_(*x*) = *θ*(*µ* − *x*), where 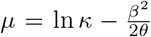, and 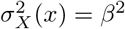. Thus, written in SDE form the diffusion *X*_*t*_ becomes

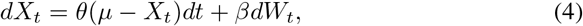

which is a special OU process, one where the mean is a function of both the strength of return to *µ*, given by *θ*, and the scaling of the environmental variance of the original process. Conversely, if we start with the OU process defined in eq. 4 and transform it using *y* = *g*(*x*) = *e^x^*, then we retrieve the Gompertz diffusion with environmental stochasticity (eq. 3). These results suggest that Palacios and Minin [1]’s transformation (eq. 1), when applied to an OU process, should result in a population growth diffusion under density-dependence and either environmental stochasticity, demographic stochasticity or both. This Itô transformation is explored in the next section.

### 2.3. Palacios and Minin’s prior on N_e_(t) as an SDE model

Palacios and Minin’s mathematical transformation of a GP in eq. 1 can be simply written as

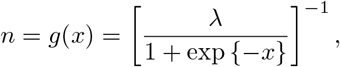

where *x* is playing the role of the GP and *n* is the effective population size. Naming the transformation as *g*(*x*) is a conventional practice in stochastic calculus and is useful to apply Ito’s transformation. Noting that *g*(*x*) can also be written as 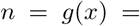 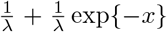, then the transformation of the OU process in eq. 4 is obtained by computing 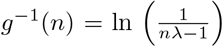 and applying Itô’s formulae to get the infinitesimal mean and variance of the *N*_*e*_(*t*) process. The infinitesimal mean is

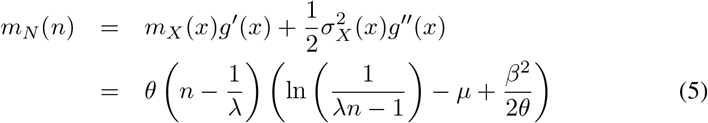

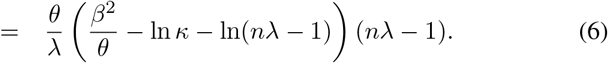

This expression is readily recognized as a translated Gompertz growth equation, one where the state space n is defined so that *n*λ − 1 > 0. The solution of the ODE 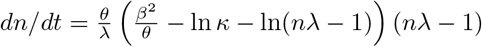 is also readily found to be

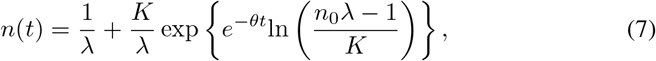

where 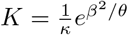 and the equilibrium state is given by 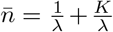. The infinitesimal variance of the resulting *N*_*e*_(*t*) diffusion is 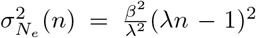. Because the process is re-scaled to have a lower bound, then this form of the infinitesimal variance is immediately recognized as the variance of a diffusion model where the quantity *n*λ − 1 displays a ‘density-dependent’, Gompertzian growth (see Figure 1) instead of logistic growth, and with added environmental variation (see Appendix 1). Thus, Palacios and Minin [1]’s prior can be cast as a fully recognizable stochastic population dynamics model, one that bears specific biological hypotheses regarding population size trajectories and their associated structure of both growth and stochasticity.

**Figure 1.**
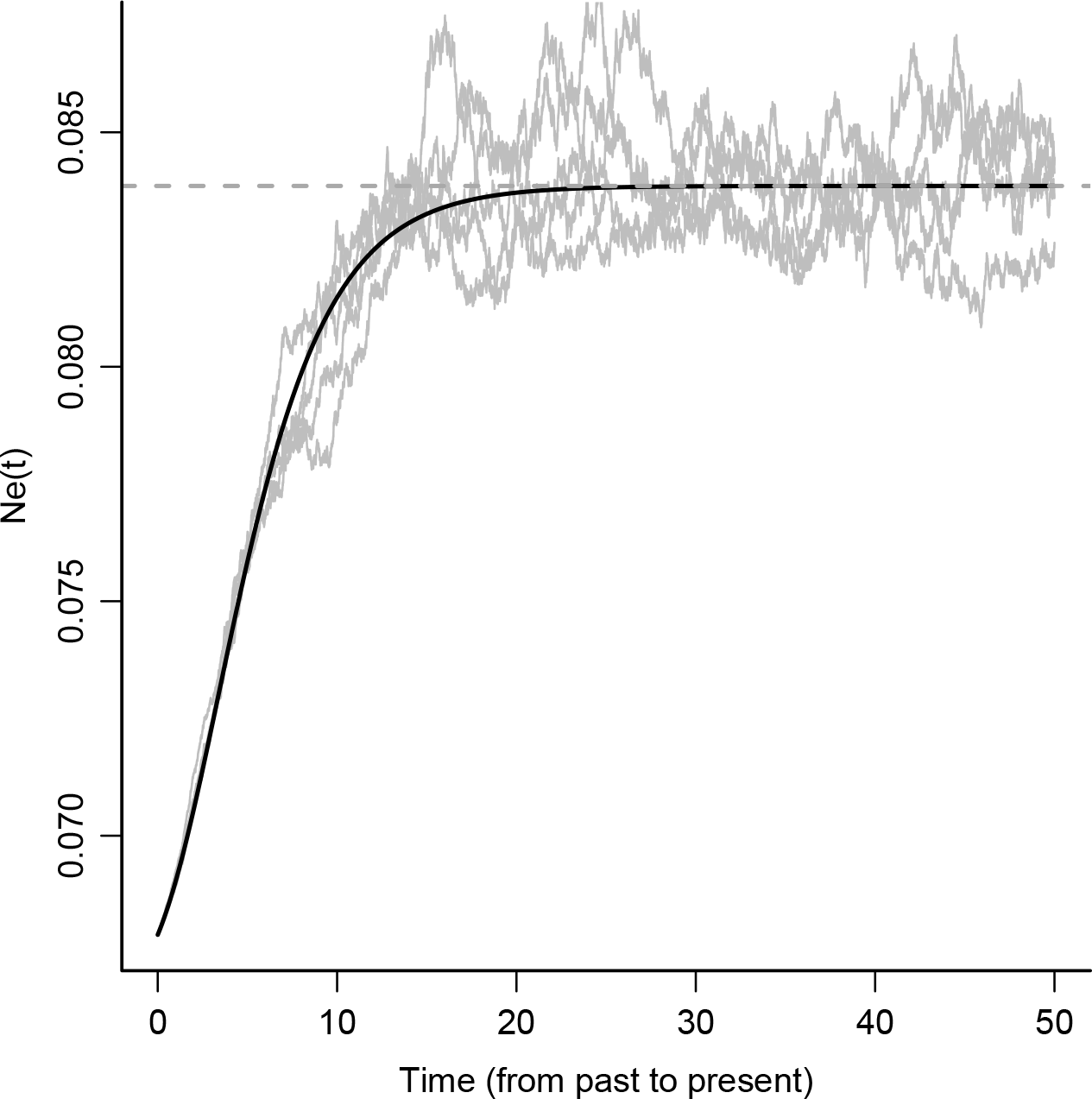
The stochastic Gompertz growth generated by the prior 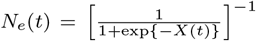, where *X*(*t*) is the OU process presented in equation 4. *N*_*e*_(*t*) is the diffusion with infinitesimal mean given by equation 6. The solid black line is the equivalent translated Gompertz trajectory (see equation 7). The horizontal dashed line corresponds to (1/*λ*) + (1/(*κ*λ)) exp(*β*^2^/*θ*). Values of the parameters used in the plot are *β*^2^ ≈ 0.002630; *κ* = 3.9140; *θ* = 0.2876;*N*_*e*_(0) = 0.0678. This plot was done by first simulating trajectories according to the OU process (equation 4), applying the prior transformation to these and then overlaying the infinitesimal mean calculation from equation 6 (as well as equation 7) as a check of calculations for the SDE transformation.

The transformed OU process according to eq. 1 has a direct connection with a class of discrete-time population dynamics models. This class of population trajectory models consists of discrete-time, density-dependent stochastic models, which Melbourne and Hastings [17] and Ferguson and Ponciano [23] show to be very flexible and accurately represent various biological systems. The connection between the OU and this class of models is possible because, as shown by Dennis and Ponciano [22], the OU process (eq. 4) has also a one-to-one transformation with the discrete-time, stochastic Gompertz model Dennis et al. [14] with environmental stochasticity, whose one-step changes are given by:

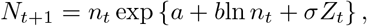

where *Z*_*t*_ ~ N(0,1) and *N*_*t*+1_ is conditional on *N*_*t*_ = *n*_*t*_. Setting *c* = *b* + 1 and *X* = ln *N*, the transition probabilities of the log-transformed Gompertz process and of the OU process coincide by setting *a* = *µ*(1 − *e^−θ^*); *c* = *e^−θ^*; *σ^2^* = (1 − *e^−2θ^*)*β*^2^/(2*θ*); *µ* = *a*/(1 − *c*) and *β*^2^ = −[2*σ*^2^ln *c*]/(1 − *c*^2^). Although the connection with both the continuous-time and the discrete-time ecological model is a rare and particular property of the Gompertz equation, it opens the door to the investigation of wether other, discrete-time ecological models can be suitably approximated by continuous processes. If carefully treated, the topic of finding equivalent, continuous-time stochastic models for these ecological processes could lead to the construction of a biologically rich (and parametric) class of priors for *N*_*e*_(*t*). To exemplify this claim, in what follows I briefly describe how thinking about these ecological models leads to *i)* a multivariate prior for a set of *p* jointly varying effective population sizes *N*_*e*_(*t*)^(*i*)^, *i* = 1, 2,…,*p* and *ii)* to a better understanding of the biological process that modulates the tempo and variance of the loss of information about past *N*_*e*_ sizes after a sudden change.

### 2.4. A multivariate prior for jointly varying effective population sizes

The stochastic Gompertz model with environmental stochasticity can readily be written in multivariate form to study the joint fluctuation of population abundances of a set of *p* species [38]. Importantly, the logarithm of this model also corresponds to a multivariate OU process, which can be subject to the same transformation used by Palacios and Minin [1] to obtain multivariate *N*_*e*_(*t*) priors. According to this Gompertz model, the set of effects of each population in each other’s rate of change, or interaction coefficients, specified as the elements of the “interactions matrix” in Ives et al. [38] formulation, determines the joint response and fluctuations of population trajectories in the face of environmental buffeting. Interestingly, this set of coefficients determines the strength of the stochasticity of the fluctuations, the rate of approach to (multivariate) stationary behavior and the speed of return to stationarity following an external perturbation of population sizes. Ives et al. [38] go as far as eliciting novel “stochastic stability” measures for multivariate time series of population trajectories. These stochastic stability measures and the corresponding multivariate prior for jointly fluctuating effective population sizes could find many applications in the study of joint gene genealogies.

### 2.5. A stochastic prior for N_e_ with a change point

The topic of parameter estimation for stochastic models with a change point is a topic that has been extensively treated in the statistical literature (see Ponciano et al (submitted) and citations therein). However, thinking of the OU-generated prior for *N*_*e*_(*t*) as a population dynamics model brings novel understanding of the nature of the the biological processes behind the speed of change, the means, variances and covariances of a stochastic *N*_*e*_(*t*) process undergoing a change. Such understanding opens the door to novel avenues of including meaningful biological insights regarding changes in *N*_*e*_.

Consider the OU process in equation 4 undergoing a change in all of its parameters (*e.g.* from *θ*_1_ to *θ*_2_, *µ*_1_ to *µ*_2_ etc…) at some point in time. If, for instance, a drop in the stationary mean of the process occurs (see figure 2), then the statistical properties of the transitional process are expressed as a function of both, the OU parameters before and after the change. In particular, it is well known that the expected value of the transitional process is a weighted average of the stationary mean before the change and the stationary mean predicted by the new set of OU parameters [39]. As time increases, the weight of the pre-change point stationary mean decreases and that of the post-change point process increases (see Figure 2). The weight that controls the loss of importance of the pre-change parameter values is 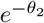. On its own, this result bears little biological significance but upon transformation of the OU process to a stochastic Gompertz prior, it is readily seen that the weight parameter controlling the loss of relevance of the past parametrization (or history), turns out to be the post-change point strength of density-dependence, 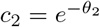 (see legend in Figure 2).

**Figure 2.**
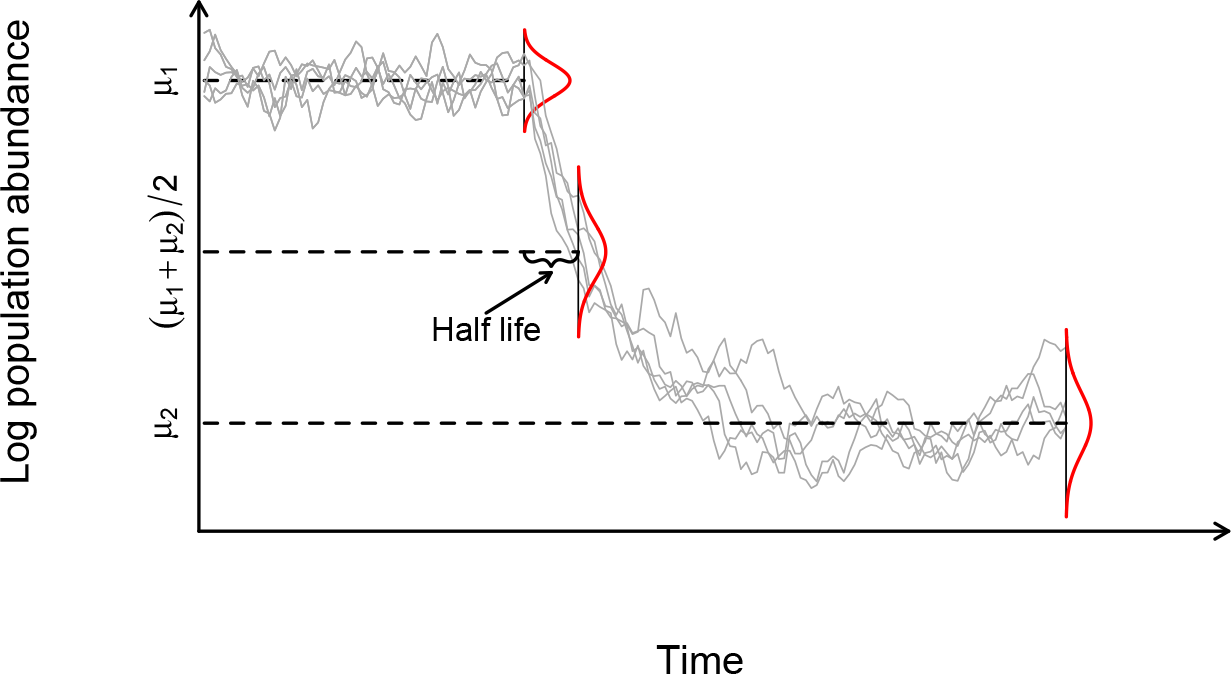
The stochastic Gompertz change-point process and the half life of the change in the mean of the dynamical process, 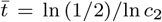. Plotted are 5 realizations of the Stochastic Gompertz model of population abundance under a change-point process. Dotted lines mark the process mean before the breakpoint (*µ*_1_), the mean after the breakpoint (*µ*_2_), and the arithmetic average of both means ( (*µ*_1_ + *µ*_2_)/2). The time at which such arithmetic average is reached is the half life of the process, 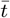. As noted in the text, 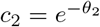 corresponds to the post-change point strength of density dependence.

The realization that Palacios and Minin [1]’s prior can be thought as a population dynamics model opens the door to biologically relevant ways to elicit priors for changes in *N*_*e*_, but defining these require analytical work not done to date (but see Ponciano et al, submitted). In statistical time series analyses and computer intensive calculations, it is customary to specify the form of the time series process with a change point as a partitioned vector. The value of the mean and variances and covariances pre and post change point are then immediately available numerically for their use in posterior calculations. At that point, all biological interpretation is lost in the computer intensive calculations. However, analytical calculations and careful simplifications show that the variances and covariances of the stochastic Gompertz SDE model similar to the *N*_*e*_(*t*) SDE shown in equation 6 can be reduced to readily interpretable variance components that are also modulated by the post-change strength of density dependence (Ponciano et al, submitted).

## 3. Discussion

In this note I show that Palacios and Minin [1]’s prior for *N*_*e*_(*t*), when obtained as a transformation of an OU process, belongs to a class of well-known stochastic population dynamics models and as such, is well rooted in a family of ecological parametric models. In ecology, the different variance scalings brought about by either type of stochasticity have enabled the separation of the contribution of environmental and demographic processes to the rate of change of the population size. If we adopt the cautious interpretation of the effective population size as a measure of the relative genetic diversity [see 1], then using the population dynamics SDE models presented here would allow the specification of different hypotheses regarding the nature of the variance components of the rate of change of genetic diversity.

Palacios and Minin [1]’s prior is indeed general because any GP, Markovian or not, can be substituted in their transformation. Thus, the nonparametric labeling of the inference they propose remains valid as well as generally applicable. However, without prior information or guidance as to what constitutes a good candidate GP, practitioners are left with a plethora of possibilities. One possibility is however, to posit as priors for *N*_*e*_(*t*) an array of models exhibiting all different combinations of the type of density dependence and structure of stochasticity in its growth rate (demographic, environmental or both).

Besides opening the door to new, obvious candidate priors (like the flexible theta-logistic model with both environmental and demographic variabilities), the results presented here also suggest novel approaches with applications in both gene genealogies inference and stochastic population dynamics modeling in ecology [23]. One of them is the formulation of a multivariate prior for jointly varying effective population sizes. Another one is the formulation of a prior for *N*_*e*_(*t*) with change points, where the strength of the changes is modulated by a parameter, which in the population dynamics world represents the strength of density-dependence, or the strength of autoregulation of the state variable of interest. In Ecology, this auto-regulation is given by intra-specific competition. The equivalent genealogical interpretation of this parameter would be that parameter which self-regulates the measure of genetic diversity.

The work presented here also reveals a first-principles justification of using the OU process to model the evolution of a quantitative trait. This process has been extensively used to model the evolution of a quantitative trait, or as it is often the case, the *logarithm* of a quantitative trait Butler and King [39], Pennell and Harmon [40]. Considerable amount of work is devoted to improve and expand this model capabilities with phenomenological modifications to the original process [*e.g* 41], and aiming to understand its benefits and limitations. It is in that respect that the connection presented here between Palacios and Minin [1]’s transformation of the OU process and stochastic population dynamics brings novel understanding into the evolution of a quantitative trait: because exponentiating an OU process results in a stochastic Gompertz model with environmental noise (stochasticity) and no demographic stochasticity, by modeling the logarithm of a trait with the OU process, a biologist is in fact hypothesizing that the trait, in its original scale grows over evolutionary time in a Gompertz-like manner (*i.e.*, ‘size-dependent’) and with a random, epochal rate of change. Furthermore, by excluding demographic stochasticity this model is stating that the total trait size is composed of equal, non-random partitions. Including ‘demographic’ stochasticity would amount to hypothesizing that the trait is composed by a sum of random, unequal contributions at any point in time.

The Itô transformation of the stochastic Gompertz model with environmental stochasticity also reveals why it is difficult to tease apart the parameter estimates *θ, µ, β*^2^ of the OU process, when used in contexts like modeling the evolution of a trait [42]. The parameter *µ* in eq. 4 is in fact, itself a function of the other two parameters and the carrying capacity *κ* of the process in its original scale. If what is being modeled is the logarithm of a quantitative trait, then the carrying capacity denotes a limiting size or constraint of the value of the trait. In fact, thinking of the branching process formulation, and because the model has no demographic stochasticity, this model is implicitly hypothesizing that a trait as a whole is conformed by the sum of individual contributions that are identical in size, and eventually converge to a given biological constrained size. A model for the evolution of a trait that includes both the analogous of demographic stochasticity and environmental stochasticity is thus, easily conceivable as a reasonable alternative to the OU process. Other forms of ‘density-dependence’ besides Gompertz can be considered too.

It could be argued that simple logistic-type models are themselves phenomenological descriptions of population growth dynamics, and thus that, the SDE Gompertz model with environmental stochasticity that results from the transformation (eq. 1) is itself phenomenological and hence, non-parametric. However, a full body of research in mathematical biosciences exists illustrating “first-principles” derivations of logistic-type models [see for instance 43].

Using well-known concepts of stochastic population dynamics, here I demonstrate that in fact, Palacios and Minin’s GP model is a special case of a population growth model with density dependence and environmental stochasticity. One of the main advantages of the Bayesian approach is the ability to include meaningful *a priori* information to conduct inference. However, eliciting priors is by far one of the most challenging problems that practitioners in population genetics and ecology face. It is in that sense that I hope that the parametric interpretation brought about by this contribution proves to be a useful, constructive critique. Finally, although the OU process has been shown to be insufficient and a current need of new parametric models has been expressed [40], its connection with stochastic, logistic-like growth pointed in this note opens the door to a plethora of other models for the evolution of a trait. Thus, Palacios and Minin [1]’s transformation has deep implications for the advancement of other areas of modeling in evolution.

## Acknowledgements

I thank professor Erkan Buzbas for leading the organization of this special issue in memory of Paul Joyce and for his critical insights as both, a reviewer and an Associate Editor. I specially thank professors Vladimir Minin, Bruce Rannala, Cécile Ané, Brian Dennis, Mark Taper and Robert D. Holt for their useful insights provided after reading early versions of this note. This work was supported by the National Institute of General Medical Sciences of the National Institutes of Health, under grant number R01GM103604 to the author.

## AppendixA. AppendixA.1. Diffusion approximation of an ecological density-dependent branching process in random environments

Let *Z*_*k*_ be the total population size at time *k* and *B*_*i*_ be the (density-dependent) offspring distribution such that *p*_*j*_(*z*) = *P*(*B*_*i*_ = *j*|*Z*_*k*_ = *z*). According to the standard definition of a branching process, we let the population size in generation *n* + 1 be given by 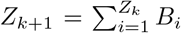. Let also denote the conditional mean and variance of the offspring distribution as *E*[*B*_*i*_|*Z*_*k*_ = *z*] = *h*(*z*) and *V*[*B*_*i*_|*Z*_*k*_ = *z*] = *v*(*z*) respectively. The idea behind the diffusion approximation is to scale both the process by some reference population size and then time. Once both the process and the original time unit have been re-scaled, in order for the diffusion approximation to hold, the first and second moment of a small increment in the process have to converge to continuous functions of the scaled process and the new time scale (see [29] eqs. 1.2 and 1.3 page 159 and eqs. 1.21 and 1.22 page 169). These two continuous functions are then the infinitesimal mean and variance of the diffusion process. The first and second moment of a one-step change in population size for the branching process above is given by:

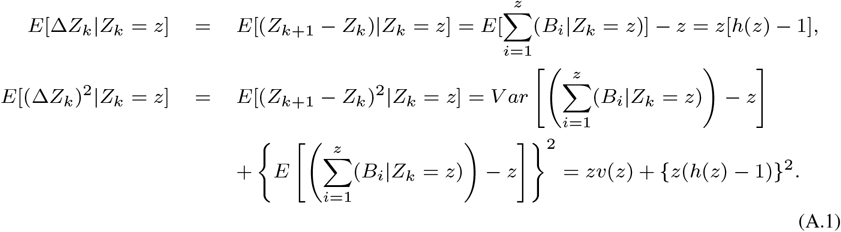

Once these moments are computed, we accelerate time by *N* (*i.e.* so that *N* generations of the original process occur in one new unit of time) and scale the state by a factor 1/*N*. Recall that the scaled process is denoted as 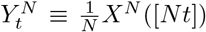, where [*Nt*] is the smallest integer close to *Nt*. With the process re-scaled, in order to find the infinitesimal mean and variance of the approximating diffusion, if it exists, we compute the infinitesimal moments and limits presented as conditions *i-iii* in the main text. For the first condition we get that

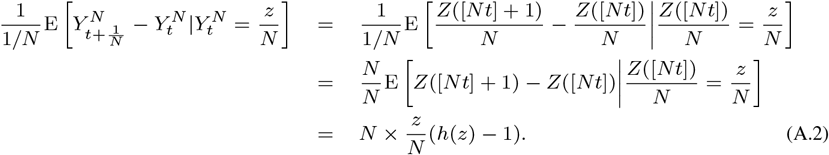

[3, 4, 6] all require that small changes in the scaled process occur in small time increments (i.e. that the offspring mean is close to replacement) because they set 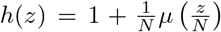. Thus, the deviations from perfect replacement are of the order 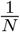. Their function *µ*(*x*), on the other hand, takes the form of the per capita growth rate given by any deterministic, single species ODE model with density dependence. Assume, for instance, that population growth conforms to the Gompertz equation shown in the main text. Then *µ*(*x*) = *θx*[log *κ* − log *x*]. Substituting this expression for the rescaled offspring mean in eq. A.2 and simplifying we get that 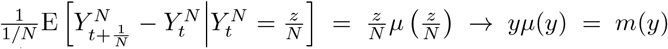 as *N* → ∞ and *z/N* → *y*. Thus, re-scaled first difference moments converge to a continuous function of y, the infinitesimal mean *m*(*y*) of the diffusion. Under the hypotheses imposed by the form of *h*(*z*), the infinitesimal mean of the diffusion can be made equal to the deterministic trend of any single species ODE model. This fact opens the door to the possibility of specifying a wide array of biological hypotheses in the form of logistic-like population growth. The second condition becomes

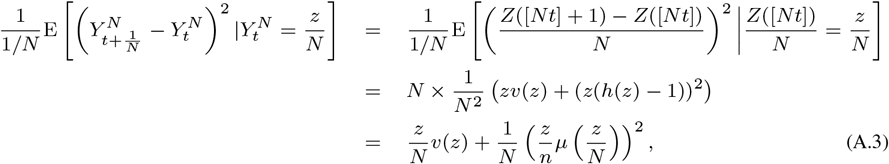

which converges to *yd*(*y*) as *N* → ∞ and *z/N* → *y*, where following [3, 4] and [6] we denote *v*(*z*) with the unspecified function of the scaled process, 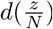. Thus, the infinitesimal variance of the diffusion is *σ*^2^(*y*) = *yd*(*y*). Therefore, both the first and the second moment converge to finite functions of *y* and *t*. The simplest assumption regarding the general function *d*(*y*) is to make it equal to a constant, say *α* > 0, which we will adopt in what follows for simplicity. [44] formally proved the convergence to a diffusion of a BP so defined. Then, the infinitesimal variance will be written as *yα*. Besides these two conditions, it is necessary for higher order moments to be negligible, *i.e.* that

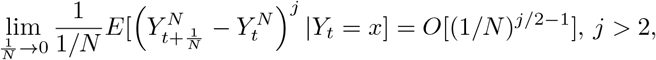

which has been done formally elsewhere [3, 4, 44]. Adding environmental stochasticity amounts to adding random fluctuations to the mean of the offspring distribution. Operationally, this is achieved by adding an *iid* process denoting random environmental fluctuations at time *k*. We denote this process with *W*_*k*_, and let *E*(*W*_*k*_) = 0 and *V*(*W*_*k*_) = 1, and assume that the *W*_*k*_ are independent from the *Z*_*m*_, *m* < *k*. Then, the probabilities of the offspring distribution are defined as

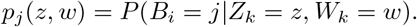

Again, *h*(*z, w*) and *v*(*z, w*) are the conditional mean and variance of the offspring distribution. To get the marginal mean and variance, using conditional expectation we average the mean and the variance of the offspring distribution over the environmental process. For the variance, authors to date have written 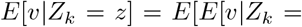 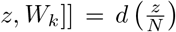. This is the expected value of the variance of the offspring distribution over the environmental process. This is then, by definition, the average demographic variance. As for the mean, adding the environmental fluctuation results in its conditional form being written as 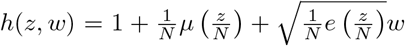, where the fluctuations due to the environment are of order 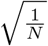 (a sum of a large number of *iid* random variables) and *e*(․) is a general function of the scaled process. As with the function *d*(․) above, the simplest assumption is to set *e*(*z*/*N*) equal to a constant, say. The diffusion approximation of a BPRE has then been found by computing the infinitesimal mean and variance of the re-scaled process as in eq. A.2 and A.3. For the first moment, and averaging over the environmental process we get that:

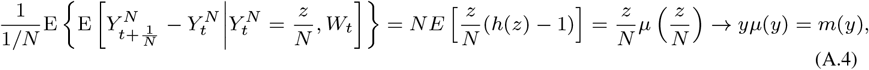

whereas for the second moment

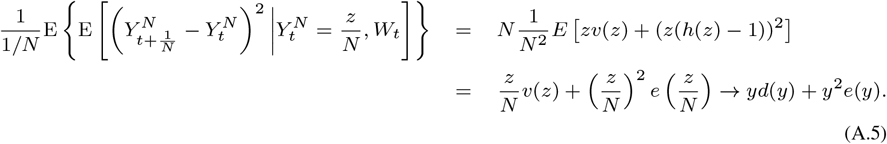

For simplicity, we assume that the functions *d*(*x*) and *e*(*x*) are constants equal to *α* and *β*^2^ respectively. Authors that have adopted this constance assumption, like [6] have implicitly stated that only the mean of the offspring distribution is affected by the environmental process. This assumption, however, has been recently been lifted by Ferguson and Ponciano [24], who also showed under which conditions the extinction probabilities predicted by the model with a constance assumption are meaningfuly under or over estimated. Finally, note that in the equation for the environmental variance, *yα* is the expected value of the variance of the offspring distribution: this term specifies on average, how much does the offspring distribution varies. The term *y*^2^*β*^2^ in turn represents the variance of the expected value of the offspring distribution: it quantifies how much does the mean of the offspring distribution changes over time.

